# Exploring the activation mechanism of the Mic60 subcomplex governing mitochondrial crista junctions

**DOI:** 10.64898/2026.10.01.755917

**Authors:** Evangelia Nathanail, Max Ruwolt, Gillian Dornan, Wiebke Y. Mikalo, Debora Elzha Aprillia Liu, Christina Schiel, Altan Tekgül, Abigail Hays Lewis, Karina von der Malsburg, Tobias Bock-Bierbaum, Volker Haucke, Séverine Kunz, Martin van der Laan, Fan Liu, Oliver Daumke

## Abstract

Crista junctions (CJs) are narrow infoldings of the inner mitochondrial membrane, serving as entry points into the cristae microenvironment. The mitochondrial contact site and cristae organizing system (MICOS) complex is a highly conserved protein machinery controlling CJ biogenesis and architecture, but its assembly mechanism driving CJ formation has remained unclear. By determining a crystal structure of a *Chaetomium thermophilum* Mic60-Mic19 complex and combining it with modelling and mutagenesis studies, we identify an extended helical region (EHR) of the Mic60 mitofilin domain and a further Mic19 helix as crucial elements of complex formation. *In vitro* cross-linking data and hydrogen-deuterium exchange mass spectrometry resolve domain assembly and conformational dynamics of the Mic60-Mic19 subcomplex and point to the EHR as a molecular switch controlling Mic60 transition from a closed dimer to a Mic19-bound hetero-octamer. Our data provide a structural model for Mic60 activation and therefore MICOS assembly as a prerequisite for crista junction formation.

## Introduction

Mitochondria are eukaryotic organelles with numerous essential cellular functions, including energy conversion, the regulation of apoptosis, fatty acid and amino acid metabolism (Kondadi & Reichert, 2024; Suomalainen & Nunnari, 2024; Tábara et al., 2025). Their multifaceted roles are reflected in their diverse network, which varies in its membrane architecture and morphology depending on organism, tissue, and energetic state (Quintana-Cabrera & Scorrano, 2023; Ryu et al., 2024). Mitochondria are enclosed by two membranes, the outer mitochondrial membrane (OMM) and the inner mitochondrial membrane (IMM), with the space in between them termed the intermembrane space (IMS) (Palade, 1952; Sjostrand, 1953). The IMM possesses invaginations, termed cristae, that serve as microenvironments of the IMS and the IMM, extending into the matrix (Mannella, 2006). They house the oxidative phosphorylation machinery (Davies et al., 2011; Wurm & Jakobs, 2006; Zick et al., 2009) and are optimized for chemiosmotic coupling of electron flux, proton pumping, and mechano-chemical ATP synthesis through their unique membrane architecture as well as lipid and protein composition (Ikon & Ryan, 2017; Mühleip et al., 2023; Venkatraman et al., 2023; Zheng et al., 2024).

The distinct crista microenvironment is maintained by the crista junction (CJ), which acts as a selective barrier between cristae and the IMS, regulating the passage of proteins, metabolites, and ions (Frey & Mannella, 2000; Nathanail et al., 2026; Rampelt et al., 2017; Wolf et al., 2019). The mitochondrial contact site and cristae organizing system (MICOS) complex is a conserved multi-subunit protein complex that localizes to CJs and plays a crucial role in their generation and structural stability (reviewed in Daumke & van der Laan, 2025; Harner et al., 2011; Hoppins et al., 2011; von der Malsburg et al., 2011). MICOS subunits are organized into the Mic10 subcomplex and the Mic60 subcomplex. The Mic12 subunit was described to bridge the two subcomplexes in yeast (Anand et al., 2016; Guarani et al., 2015; John et al., 2005; Mun et al., 2010; Naha et al., 2024). Deletion of Mic60 or Mic10 nearly abolishes CJ formation, with cristae remaining mostly disconnected from the IMS in the matrix (Bohnert et al., 2012; Harner et al., 2011; Hoppins et al., 2011; von der Malsburg et al., 2011). The Mic10 subcomplex, consisting of Mic10, Mic26 and Mic27 is composed of largely membrane-embedded wedge-shaped subunits, believed to form a scaffold that stabilizes CJ curvature (Barbot et al., 2015; Bohnert et al., 2015; Brown et al., 2026; Meena et al., 2025; Stephan et al., 2024). In contrast, the Mic60 subcomplex (consisting of Mic60, Mic19 and Mic25 in vertebrates) is anchored to the IMM only by the transmembrane helix (TM) of Mic60 and is mostly exposed to the IMS (Daumke & van der Laan, 2025). Apart from its role in shaping CJs, the Mic60 subcomplex forms the mitochondrial intermembrane space bridging complex (MIB) by binding to the OMM-localized sorting and assembly machinery (SAM) complex, implicating a function in mitochondrial protein trafficking and import (Korner et al., 2012; Tang et al., 2020; Xie et al., 2007; Zerbes et al., 2012). In yeast, the Mic60 and Mic10 subcomplexes localize to CJs independently of each other (Friedman et al., 2015; Tirrell et al., 2020). Their assembly is regulated by factors such as cardiolipin (Rampelt et al., 2018) and, in human cells, optical atrophy 1 (OPA1) (Stephan et al., 2020), which are both closely tied to cristae shape. Establishing the molecular assembly mechanism of the MICOS subcomplexes is therefore essential for understanding cristae biogenesis and mitochondrial architecture.

Biochemical and structural studies of the Mic60-Mic19 complex from thermophilic fungal orthologues showed that the nanomolar-affinity interaction of the two proteins is driven by the Mic60 C-terminal mitofilin domain and the Mic19 Coiled-coil-helix-coiled-coil-helix (CHCH) domain, while further regions in Mic60 and Mic19 may contribute to the assembly (Bock-Bierbaum et al., 2022; Hessenberger et al., 2017). The mitofilin domain includes two peripheral lipid binding sites (LBS) called LBS1 and LBS2 (Hessenberger et al., 2017). In the published structure of the mitofilin-CHCH complex, the mitofilin domain forms a domain swapped dimer (Bock-Bierbaum et al., 2022). Disruption of this mitofilin-CHCH domain interaction abolishes complex formation and impairs CJ assembly in a yeast model.

Upon Mic19 binding, Mic60 transitions from a dimeric, autoinhibited state to a higher-order oligomeric state (Bock-Bierbaum et al., 2022; Hessenberger et al., 2017). Central to the formation of the assembled hetero-octamer is the coiled-coil (CC) domain of Mic60, which forms a stable antiparallel dimer and further assembles into a cross-shaped tetramer via a conserved interface (Bock-Bierbaum et al., 2022). The fully formed Mic60-Mic19 subcomplex was proposed to act as a mechanical strut, spanning across the CJ (Bock-Bierbaum et al., 2022), while modelling studies suggested that the human subcomplex acts as a diffusion barrier by physically occupying the CJ space (Nathanail et al., 2026). However, little is known about the mechanisms governing Mic60-Mic19 complex formation. Capturing dynamic intermediate states and identifying key hinge regions would advance our understanding of MICOS assembly and its role in CJ biogenesis.

Here, we investigated Mic60-Mic19 subcomplex formation from the model organism *Chaetomium thermophilum* using a combination of biophysical, biochemical and structural experiments. We revealed the crystal structure of an N-terminally extended mitofilin domain of Mic60 bound to the CHCH domain of Mic19 and identified an extended helical region (EHR) of Mic60 as an essential determinant of complex formation with Mic19 in yeast. AlphaFold-guided structural modelling identified an additional interaction site of the Mic60 coiled-coil domain with a predicted α-helix in Mic19, which we biochemically validated. To probe the role of the EHR in Mic60-Mic19 assembly and to characterize the conformational dynamics associated with complex formation, we performed *in vitro* cross-linking mass spectrometry and hydrogen-deuterium exchange mass spectrometry. Our data lead us to propose a structural model of how Mic60 transitions from a closed homo-dimer to an open, extended Mic60-Mic19 hetero-octamer, centring around the EHR as a structural switch.

## Results

### Identification of the EHR in the extended Mic60 mitofilin domain

Mic60-Mic19 complex formation is driven by the mitofilin-CHCH interaction, but it involves also regions beyond this domain interface, since the highest binding affinity between the two proteins is observed with full-length, not truncation constructs (Hessenberger et al., 2017). We sought to characterize these additional interactions in molecular detail by recombinantly expressing and purifying N-terminally extended constructs of the mitofilin domain of Mic60 and the CHCH domains of Mic19 (Figure 1A, Supplementary Figure 1A-C). In isothermal titration calorimetry (ITC) experiments, the extended mitofilin construct bound full-length Mic19 and the isolated CHCH domain with high affinity (Supplementary Figure 1D). Consequently, the extended mitofilin and CHCH constructs co-eluted in gel filtration, forming a heterodimer of 34 kDa, as detected by right-angle light scattering (SEC-RALS) (Figure 1B). In order to gain molecular insight into this interaction, crystals of the extended mitofilin domain in complex with the CHCH domain were grown and diffracted to 2.11 Å. The phase problem was solved by molecular replacement using an AlphaFold2 prediction of the purified constructs as a search model (see Materials and Methods, Supplementary Figure 1E) and the model was refined to an R_work_/R_free_ of 23.1%/27.8% (see Table 1 for data collection and refinement statistics).

**Figure 1:**
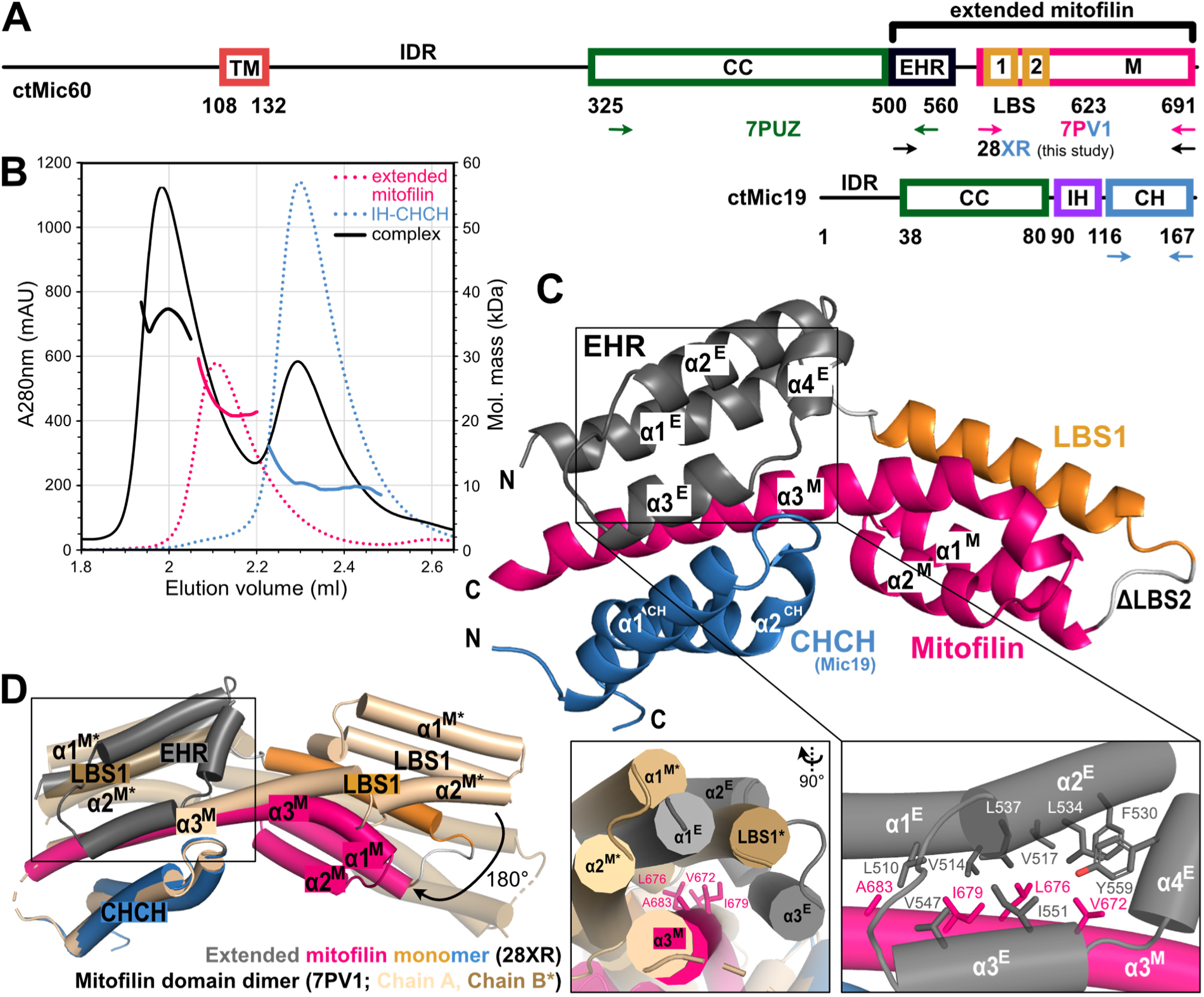
The mitofilin domain of Mic60 harbours an extended helical region (EHR). **A** Schematic domain architecture of Mic60 and Mic19, with residue numbers annotated below the domains. Intrinsically disordered regions are annotated as IDR, transmembrane helix (TM) is coloured in red, coiled-coil (CC) in forest green, extended helical region (EHR) in dark grey, lipid binding sites 1 and 2 (LBS) in orange, mitofilin domain (M) in hot pink, interface helix (IH) in purple, Coiled-coil-helix-coiled-coil-helix (CHCH) domain in skyblue. Solved crystal structures and their domain boundaries in *ct*Mic60 are annotated by arrows and their PDB ID (ltMic60 CC 7PUZ in green, *ct*Mic60 LBS1-mitofilin-CHCH 7PV1 in hot pink and skyblue, *ct*Mic60 extended mitofilin-CHCH 28XR in black and skyblue). **B** SEC-RALS profile showing the co-elution of the extended mitofilin-CHCH complex as a hetero-dimer in solution. **C** Cartoon representation of the structure of the extended mitofilin-CHCH complex, with a close-up view of the four-helix bundle of the EHR bound to mitofilin helix 3 (α3^M^). Domains are coloured according to A and helices are annotated and numbered accordingly (E: EHR, M: mitofilin). ΔLBS2 indicates the deletion of LBS2. **D** Superposition of the extended mitofilin crystal structure solved here and 7PV1. Note that 7PV1 is a mitofilin-CHCH fusion construct that assembles into a domain-swapped dimer (chain A coloured in wheat, chain B in sand and with helices indicated by asterisks). A close-up view of a hydrophobic patch of α3^M^ illustrates the clash between the LBS-mitofilin bundle of the second moiety of 7PV1 (sand) and the EHR of the extended mitofilin structure (grey).

Consistent with the SEC-RALS analysis (Figure 1B), the structure of the complex comprises a monomer of the extended mitofilin domain bound to a monomer of the CHCH domain (Figure 1B). A three-helix bundle was present N-terminally of the mitofilin domain that was not included in the previously crystallized mitofilin domain construct. The three-helix bundle is composed of helices α1^E^-α3^E^ and was capped on one side by the short helix α4^E^. From here, we will refer to this region as the Extended Helical Region (EHR, *ct*Mic60 aa 500-562, Figure 1C). Notably, the EHR sequence was identified before in evolutionary analyses of Mic60 orthologues as part of an ‘extended mitofilin domain’ (Benning et al., 2025).

The EHR binds through a conserved hydrophobic core to the central helix α3^M^ of the mitofilin domain, opposite of the Mic19 binding interface (Figure 1C, close-up 1, Supplementary Figure 2). Interestingly, the same hydrophobic surface of α3^M^ was found to pack against the LBS bundle of the opposite monomer in the domain-swapped mitofilin domain dimer (PDB ID: 7PV1, Figure 1D) (Bock-Bierbaum et al., 2022). When comparing the two structures, the LBS bundle undergoes a 180° rotation along the central mitofilin helix, while preserving its overall fold (Figure 1D, Supplementary Figure 3A-C). Thus, depending on the packing of α3^M^ against the LBS or EHR bundle, the mitofilin domain adopts a monomeric or a dimeric assembly.

We had previously suggested that the convex architecture of the mitofilin dimer contributes to binding the highly curved CJ neck. An electrostatic surface potential of the extended mitofilin monomer shows a vast positively charged interface that could potentially also interact with the membrane as a monomer (Supplementary Figure 3D, E). In summary, the extended mitofilin region of Mic60 can form both monomers and dimers in complex with the CHCH domain of Mic19.

### The EHR participates in and is required for Mic19 binding

The primary interface of the Mic60 mitofilin and Mic19 CHCH domain in our crystal structure was essentially identical to that of the published Mic60-Mic19 fusion construct - the corresponding regions could be superimposed with a low root mean square deviation of 0.3 Å (Figure 2A, B). However, in addition to the primary contact site, EHR helix α3^E^ interacted with the first CHCH helix α1^C^ through conserved Mic60 residues V546 and A549 (Figure 2A, Supplementary Figure 2A). These pack against Mic19 residues V129 and L132, previously described as essential for Mic60-Mic19 complex formation *in vitro* and *in vivo* (Bock-Bierbaum et al., 2022).

**Figure 2:**
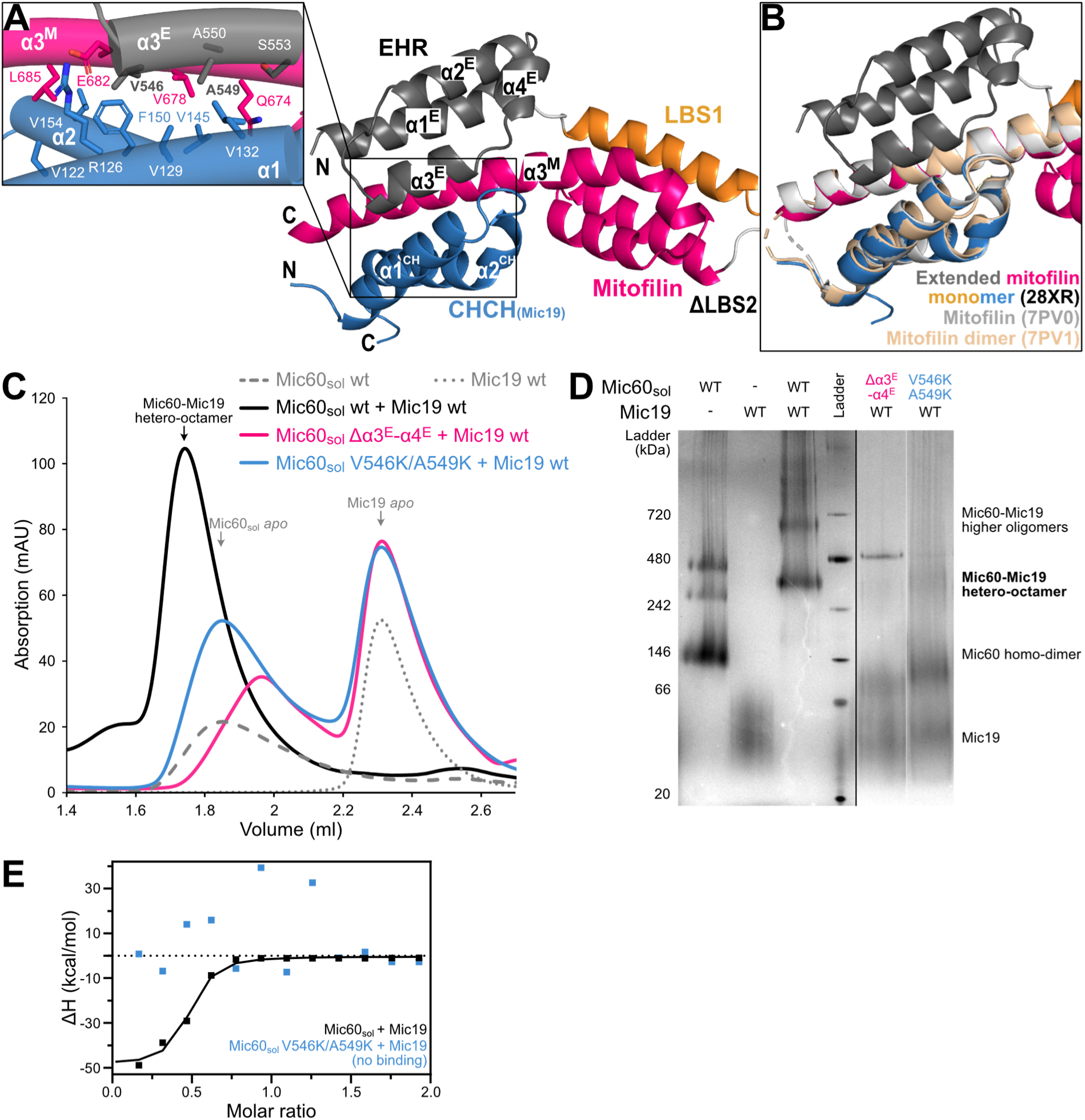
The EHR contributes to Mic19 binding *in vitro*. **A** Cartoon representation of the structure of the extended mitofilin-CHCH complex, with a close-up view of the interaction of the CHCH domain of Mic19 (blue), α3^E^ of the EHR (black) and the α3^M^ (pink). **B** Superposition of the crystal structure of the extended mitofilin domain and the previously solved structures of mitofilin-CHCH fusion constructs (PDB IDs: 7PV0 in light grey and 7PV1 in wheat). Alignment and root mean square deviation (RMSD) calculation were performed with the residues present in 7PV0 (C-terminus of α3^M^ and the CHCH domain), RMSD was 0.3 Å for 7PV0 and 0.6 Å for 7PV1. **C** Analytical SEC showing aberrant Mic60-Mic19 complex formation upon deletion of EHR helices α3^E^-α4^E^ (pink), and of the CHCH-contacting residues V546 and A549 (blue). **D** BN-PAGE gels showing aberrant Mic60-Mic19 complex formation upon modification of the EHR as shown and coloured in C. Bands are categorized on the right side of the gel as in Bock-Bierbaum et al. (2022). For Mic60_sol_ *apo* controls see Supplementary Figure 4A. **E** ITC profile of the high-affinity binding of WT Mic60_sol_ to Mic19 (black, K_D_ = 600 ± 200 nM, binding number n = 0.43 ± 0.01), and the lack of binding upon substituting Mic60 residues V546K/A549K (blue).

To dissect the contribution of the EHR to Mic60-Mic19 binding, we performed site-directed mutagenesis and analysed complex formation by analytical SEC and Blue Native polyacrylamide gel electrophoresis (BN-PAGE). In agreement with previous data, a wild-type Mic60 construct comprising the soluble Mic60 intermembrane space domain (Mic60_sol_) transitioned from a basal dimeric state to a higher order oligomeric assembly upon Mic19 binding, eluting as a complex on SEC (Figure 2C) and forming a distinct oligomeric band on BN-PAGE (Figure 2D) (Bock-Bierbaum et al., 2022; Hessenberger et al., 2017). In contrast, deletion of helices α3^E^-α4^E^ or modification of the newly identified EHR Mic60-Mic19 contact point in the V546K/A549K double variant abolished complex formation (Figure 2C) and hetero-octamer assembly (Figure 2D). While Mic60_sol_ bound with high affinity to Mic19 (K_D_ = 600 ± 200 nM) in isothermal titration calorimetry (ITC) experiments, the V546K/A549K variant did not show a binding signal, indicating that the substitutions disrupt the interaction (Figure 2E). Perturbing EHR helices α1^E^-α2^E^ through deletion or by substitution of conserved residues V514D/V517D or F530D/L534D, reduced yet did not abolish complex formation in BN-PAGE and analytical SEC (Supplementary Figure 4A, B), highlighting EHR helix α3^E^, and particularly residues V546 and A549, as vital for Mic19 binding.

To analyse the role of the new interactions for CJ formation, we employed *S. cerevisiae* as an established model system (Bock-Bierbaum et al., 2022). To this end, we re-expressed wild-type (WT) Mic60 and an EHR variant (L407K/V410K in yeast, equivalent to *ct*Mic60 V546K/A549K) in a Mic60 deficient yeast strain, in which Mic19 was fused and therefore stabilized with a C-terminal Protein-A tag (PA). Indeed, expression levels of the Mic60 variant were similar compared to WT. However, whereas WT Mic60 was efficiently pulled-down with Mic19-PA together with further MICOS subunits, the EHR variant was not, indicating that Mic60 subcomplex formation was disturbed by the amino acid exchanges (Figure 3A).

**Figure 3:**
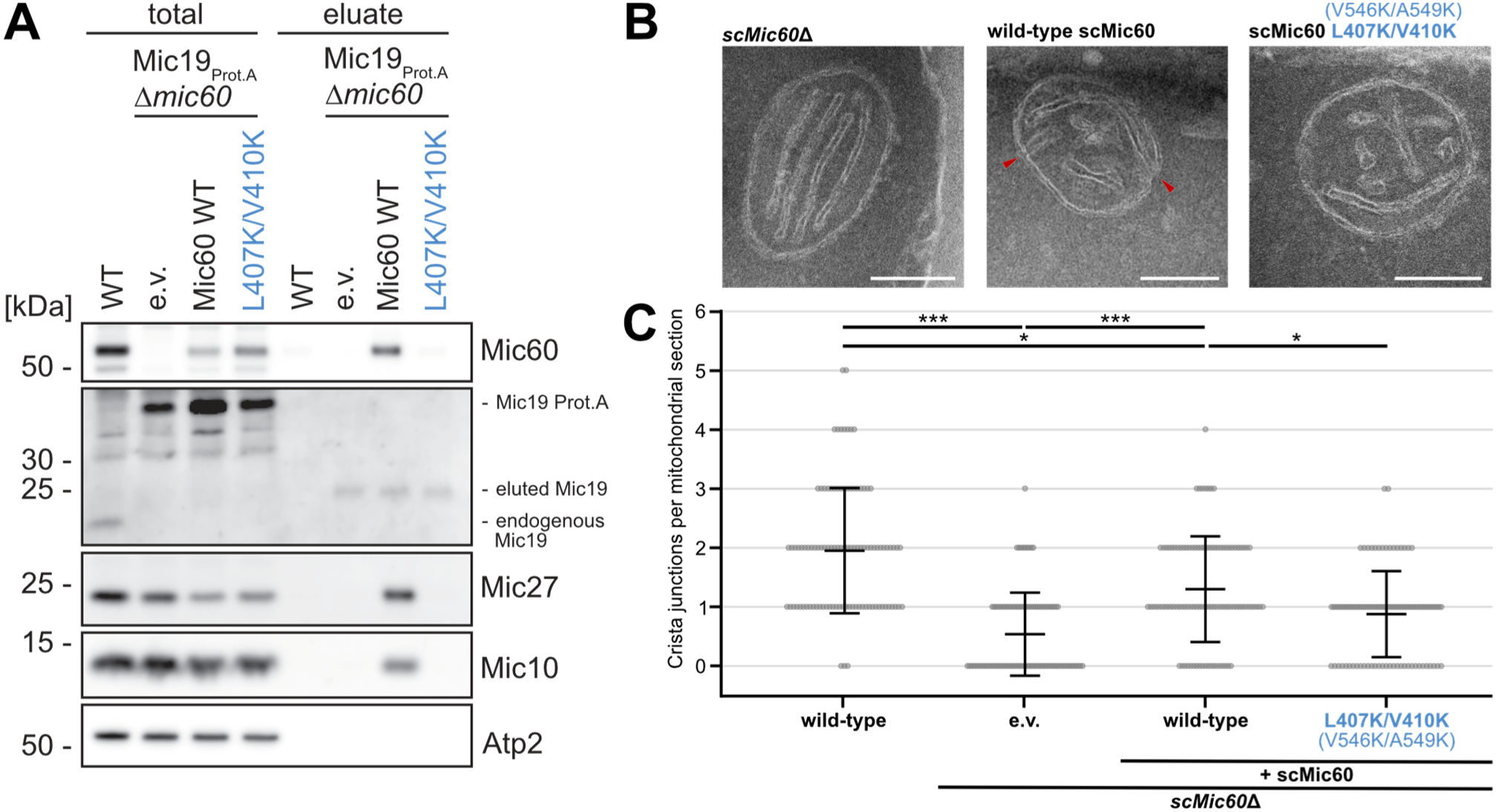
The EHR supports CJ formation in yeast cells. **A** Western blot of immunoprecipitation experiments with isolated *S. cerevisiae* mitochondria containing Mic19-Protein A fusion protein, lacking the chromosomal *MIC60* gene (*mic60*Δ) and transformed with an empty vector (e.v.) or plasmids encoding wild-type (WT) or modified Mic60 as indicated. **B** Representative electron microscopy micrographs of the mitochondria from *S. cerevisiae* cells analysed in A, cut in ultrathin sections. The numbering of the corresponding *ct*Mic60 residues is in brackets. The white bar represents 200 nm. **C** Quantification of CJs per mitochondrial section in the *S. cerevisiae* cells shown in G. *** for p < 0.001, * for 0.01 ≤ p < 0.05 (n = 100 mitochondria, showing mean and standard deviation).

We next analysed the consequence of the substitution for CJ formation. To this end, yeast cells were fixed and ultra-thin sections prepared for EM analyses. As previously described (Bock-Bierbaum et al., 2022), deletion of Mic60 led to a loss of CJs, which could be rescued by re-expression of WT Mic60, but not of an empty vector (Supplementary Figure 5). Re-expression of Mic60 with the same double substitution in the EHR domain led only to a partial rescue of CJ formation (Figure 3B, C), indicating an important role of the Mic60 EHR - Mic19 CHCH domain interaction for the formation of CJs.

### The Interface Helix (IH) of Mic19 participates in complex formation through a secondary binding site

To obtain further information on the Mic60-Mic19 assembly, we generated a structural model of the *Chaetomium thermophilum* (*ct*) Mic60-Mic19 complex using AlphaFold2 (Figure 4A, Supplementary Figure 6A) (Jumper et al., 2021). The model strikingly recapitulated the interfaces we had observed in our crystal structure, including the primary mitofilin domain-CHCH interface and the EHR-CHCH domain interaction (Site 1). In addition, a second interaction site in the predicted complex involved a structurally uncharacterized α-helix of Mic19, here referred to as Interface Helix (IH). In the predicted model, the IH engaged with a C-terminal segment of the Mic60 CC, here referred to as the CC tail (Site 2).

**Figure 4:**
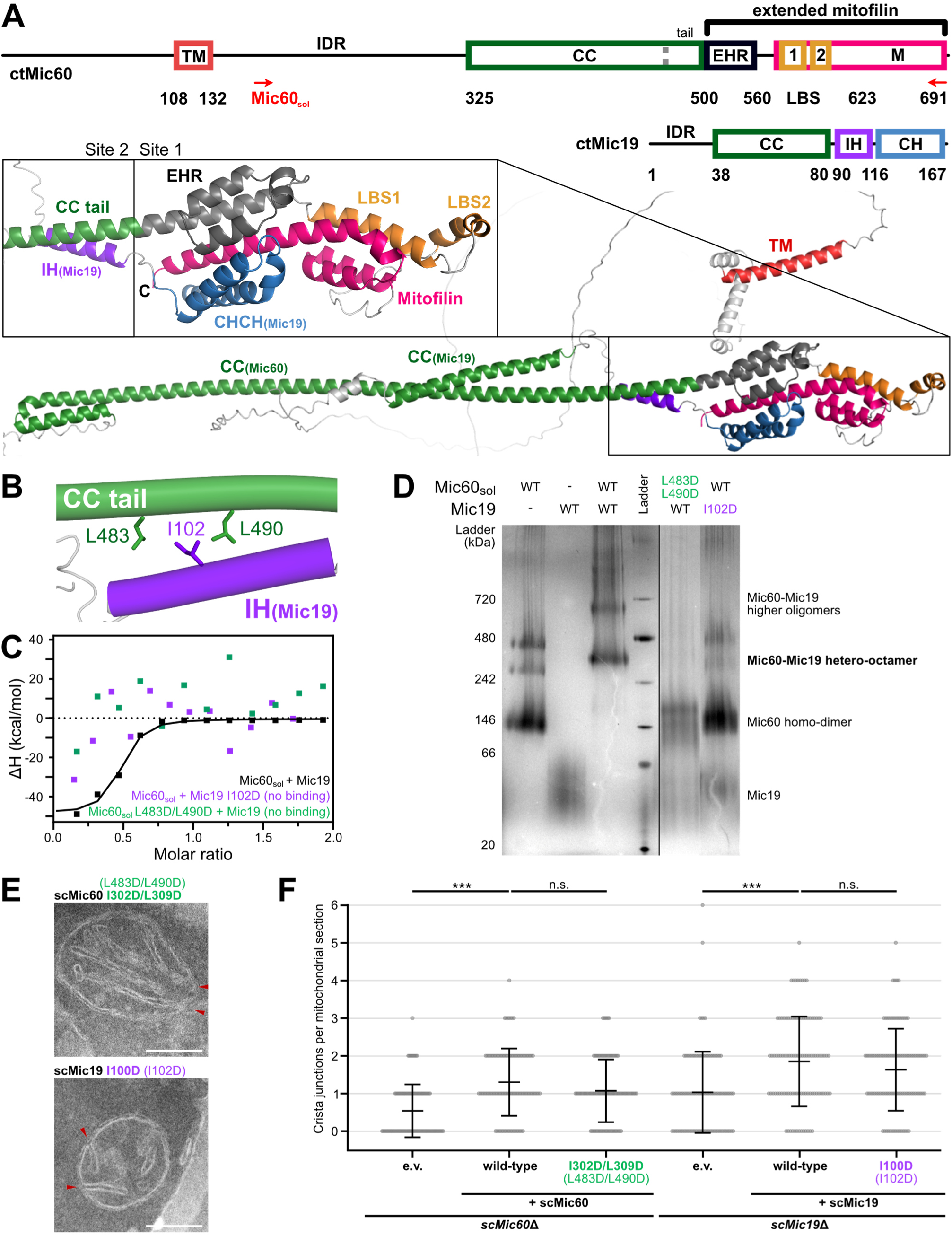
The CC of Mic60 and the IH of Mic19 form a secondary binding site that contributes to Mic60-Mic19 complex formation. **A** Schematic domain architecture of Mic60 and Mic19 as in Figure 1A. Construct Mic60_sol_ (*ct*Mic60 aa 208-691) is annotated by red arrows. Cartoon representation of the AlphaFold2 prediction of the full-length *ct*Mic60-*ct*Mic19 complex, coloured according to the domain annotations above, with a close-up view of the C-terminal interaction interface, which is separated into interaction site 1 (CHCH-mitofilin) and 2 (IH-CC tail). For AlphaFold confidence scores, see Supplementary Figure 6A. **B** Close-up view of the secondary binding site IH-CC tail. **C** ITC profile of the high-affinity binding of WT Mic60_sol_ to Mic19 (black, as seen in Figure 2E), and the lack of binding upon modification of Mic60 L483D/L490D (green) or Mic19 I102D (purple). **D** BN-PAGE gels showing aberrant Mic60-Mic19 complex formation upon modification of the IH-CC tail interface, as shown in C. Bands are categorized on the right side of the gel as in Bock-Bierbaum et al. (2022). **E** Representative micrographs of *S. cerevisiae* cells that contain a *sc*Mic19-Protein A fusion, lack the chromosomal *MIC60* gene (*mic60*Δ) and were transformed with an empty vector (e.v.) or with plasmids encoding wild-type or modified *sc*Mic60 as indicated. For Mic19 variant analysis cells harbor a *sc*Mic60-Protein A fusion, lack the chromosomal *MIC19* gene (*scMic19*Δ) and were transformed with an empty vector (e.v.) or with plasmids encoding wild-type or modified *sc*Mic19 as indicated. **F** Quantification of CJs per mitochondrial segment of *S. cerevisiae* cells as shown in E. For the scMic60 analysis, cells *** for p < 0.001, not significant (n.s.) for p ≥ 0.05 (n = 100 mitochondria, showing mean and standard deviation). For further representative micrographs see Figure 3B and Supplementary Figure 5.

We characterized the new IH-CC tail binding interface (Figure 4B) by altering conserved residues in the Mic60 CC tail (L483D/L490D) or the Mic19 IH (I102D) (Supplementary Figure 2A). By employing ITC, we observed that Mic60-Mic19 binding was abolished by the amino acid substitutions in our reconstituted system using purified proteins, indicating that the IH-CC tail interface contributes to complex formation (Figure 4C). BN-PAGE analysis confirmed these data, showing disrupted hetero-octameric complex formation for the variants, although partial hetero-tetrameric assembly was still observed in the form of a second band above the Mic60 *apo* band for the L483D/L490D variant (Figure 4D) and by the lack of excess Mic19 on analytical SEC (Supplementary Figure 4B).

To determine the role of the IH-CC tail interface in yeast cells, we re-expressed a Mic60 variant with the corresponding amino acid substitutions in the CC tail (I302D/L309D, equivalent to *ct*Mic60 L483D/L490D) or a Mic19 PA-fusion construct containing an alteration in the IH (I100D compared to I102D in *ct*Mic19) in the corresponding knockout strains. None of the exchanges affected the expression levels of the respective proteins. When probed in pull-down analyses, these substitutions did not affect Mic60 subcomplex formation (Supplementary Figure 6B). Consistently, when yeast cells were imaged by transmission electron microscopy, mitochondria with modified Mic60 or Mic19 exhibited CJs comparable in size and number to those of that harboured the respective wild-type proteins (Figure 4E, F). These experiments suggest that IH-CC tail association is not essential for Mic60 subcomplex formation in yeast and that compensatory mechanisms may still allow for the formation of CJ junctions *in vivo* despite a disturbed IH-CC tail interface.

### Unbound Mic60 forms a dimer in a closed conformation

Structural comparisons revealed that EHR helices α1^E^ and α2^E^ were already previously described in the structure of the Mic60 coiled-coil construct of *Lachancea thermotolerans* (PDB ID: 7PUZ), where they packed against the CC tail. Interestingly, the EHR residues interacting with the CC tail in the previously published structure pack against the mitofilin domain in the current structure, meaning the two binding modes of the EHR towards the coiled-coil or the mitofilin domain are mutually exclusive (Figure 5A). We therefore wondered whether the EHR may function as a molecular switch: in the absence of Mic19, the EHR could pack against the dimeric CC, while in the presence of Mic19, it may be displaced from the CC tail by the Mic19 IH and instead bind to the mitofilin domain.

**Figure 5:**
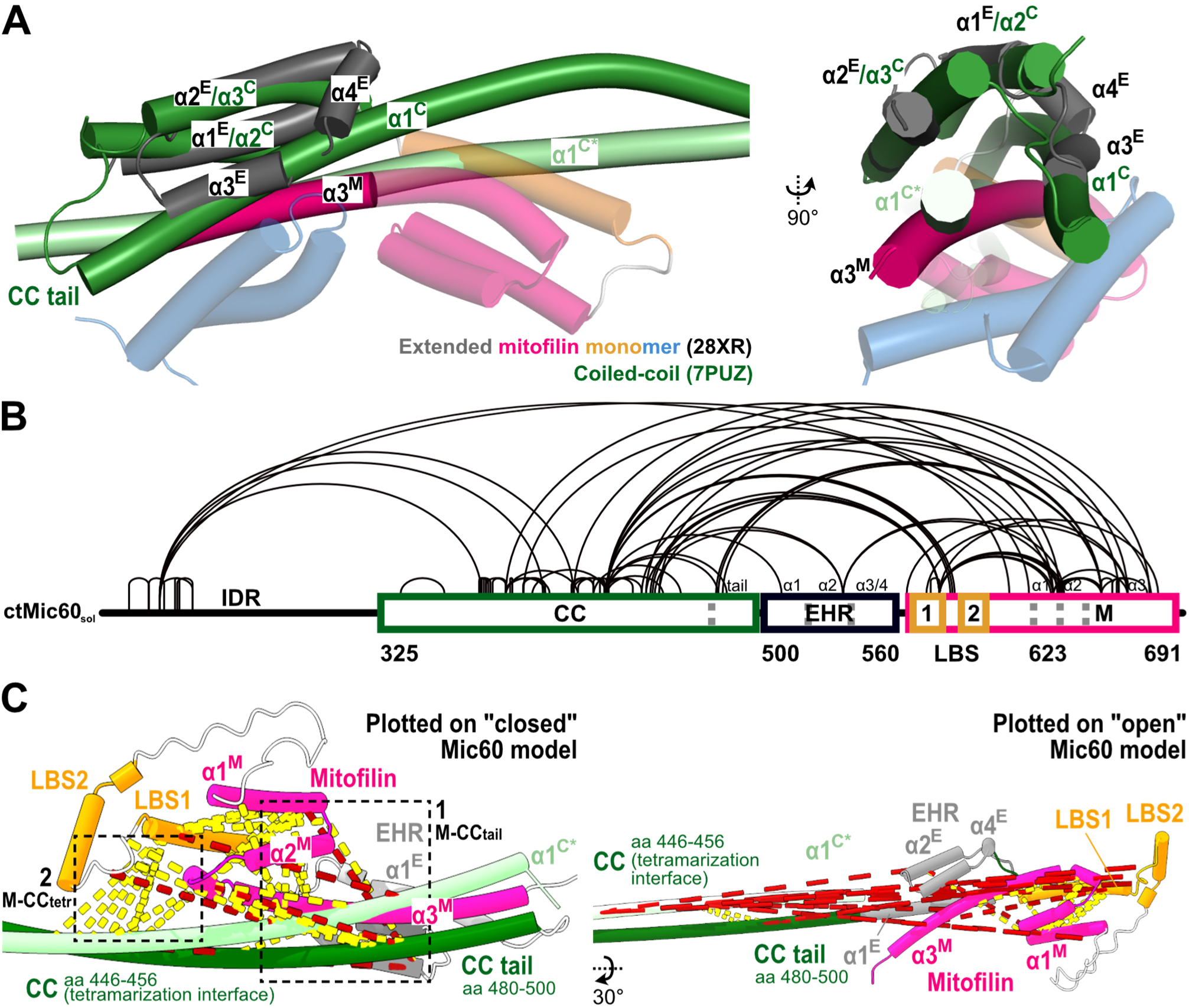
Cross-linking MS resolves the extended mitofilin domain in the Mic60 unbound state. **A** EHR helices α1^E^ and α2^E^ were resolved as part of the coiled-coil structure of ltMic60 (equivalent to α2^C^ and α3^C^ of the CC from PDB ID 7PUZ), showing here the superposition of the two structures with an RMSD calculated for the two helices at 2.1 Å. From the tetrameric assembly of the coiled-coil, only one antiparallel dimer is shown here, with chain A coloured in forest green and chain B in light green. **B** Visualisation of the SDA *apo* Mic60 cross-linking dataset (n=83) on the Mic60 sequence, generated with xiNET (Combe et al., 2015). **C** SDA cross-links from B are plotted on putative models of Mic60 in its ‘closed’ (left) and ‘open’ (right) conformations, visualized using XMAS in ChimeraX (Lagerwaard et al., 2022; Meng et al., 2023); in-range cross-links are shown in yellow and out-of-range are shown in red (cut-off: 30 Å). Boxes annotate the prevalent satisfied cross-links between the extended mitofilin domain and CC tail (1) and the extended mitofilin domain and the CC tetramerization interface (2).

To characterize these conformational transitions during complex formation, we initially aimed to study the EHR conformation in the *apo* state of Mic60. We were not able to determine a crystal or cryo-EM structure for full-length Mic60, likely reflecting its intrinsic flexibility and the appearance of several assembly forms (Figure 2D). In an alternative structural approach, we performed *in vitro* cross-linking mass spectrometry (XL-MS) using the Mic60_sol_ construct in its unbound, predominantly dimeric form. For this purpose, we employed the zero-length cross-linker Succinimidyl 4,4’-azipentanoate (SDA) (Iacobucci et al., 2018) to achieve the highest possible spatial resolution in this method from ultra-short distance restraints.

*Apo* Mic60 was characterised by numerous self-cross-links connecting the CC to the extended mitofilin region (Figure 5B). To dissect and structurally exploit these cross-links, we plotted them on two putative models of Mic60 organization, guided by the two alternative conformations of the EHR (Figure 5A, see Materials and Methods). In the first ‘closed’ model, the extended mitofilin domain was in close proximity with the CC tail. In the second model, termed the ‘open’ conformation, the EHR was dissociated from the CC tail and adopted the fold observed in the extended mitofilin region structure, bound to the CHCH domain of Mic19.

We mapped the Mic60 *apo* cross-linking data onto both putative conformations and observed a higher satisfaction rate for the “closed” model (using a 30 Å cut-off (Ser et al., 2025), Figure 5C left, Supplementary Figure 8C), meaning the detected cross-links better reflected this architecture. Firstly, numerous cross-links between the extended mitofilin domain and the CC supported their close spatial proximity (box 1). Accordingly, the majority of the SDA *apo* cross-links were out of range in the model featuring the ‘open’ conformation (Figure 5C right), highlighting that the extended mitofilin domain must be in close structural proximity to the CC in the unbound state. Secondly, the LBS-mitofilin bundle cross-linked extensively with the tetramerization interface (box 2), which supports a model in which this contact is inhibiting tetramer formation, as observed on BN-PAGE.

For the closed Mic60 dimer, we initially modelled the extended mitofilin domain as two separate entities (Supplementary Figure 7A). However, we also considered a model in which the core mitofilin domain separated from the EHR and formed a domain-swapped dimer above the CC tetramerization interface (Supplementary Figure 7B), as reported in our previous publication (Bock-Bierbaum et al., 2022). Supporting such arrangement, multiple SDA *apo* cross-links were detected directly in the mitofilin dimerization interface (Supplementary Figure 7C).

### The EHR is a switch regulating Mic60-Mic19 complex assembly

Our structural and biochemical data suggested that the EHR may play a role in the conformational changes in Mic60 and serve as a molecular switch for Mic60-Mic19 assembly. To characterize these changes, we performed SDA-based cross-linking MS on the Mic60-Mic19 complex and compared it to our Mic60 *apo* dataset.

The resulting cross-linking patterns differ markedly between the two states (Figure 6A). The extensive cross-links connecting the CC to the extended mitofilin region in the *apo* dataset were largely absent in the Mic19-bound spectra. Instead, complex formation gave rise to cross-links between the Mic60 CC tail and the Mic19 IH.

**Figure 6:**
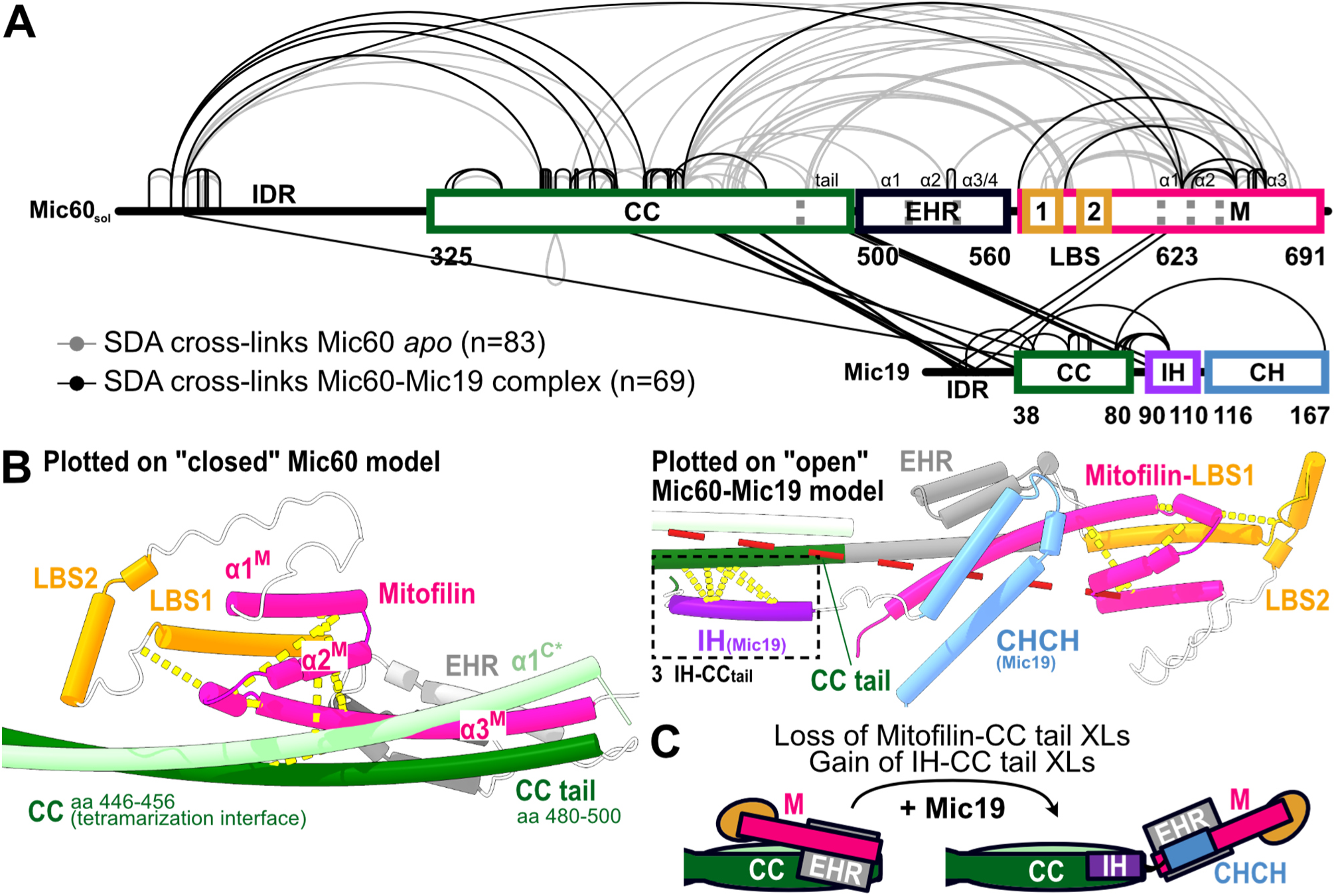
The EHR switches conformations upon Mic60-Mic19 complex formation. **A** Visualisation of the SDA cross-linking datasets on the Mic60 and Mic19 sequence, generated with xiNET (Combe et al., 2015). *Apo* Mic60 cross-links as in Figure 5B coloured in grey (n=83), Mic60-Mic19 complex cross-links coloured in black (n=69). **B** Mic60-Mic19 SDA cross-links from A are plotted on Mic60 in its putative ‘closed’ (left) and ‘open’ (right, bound to Mic19) conformations, visualized using XMAS in ChimeraX (Lagerwaard et al., 2022; Meng et al., 2023); in-range cross-links are shown in yellow and out-of-range are shown in red (cut-off: 30 Å). Box 3 indicates the cross-links between the Mic60 CC tail and the Mic19 IH. **C** Schematic representation of Mic60 transitioning from a closed to an open conformation upon Mic19 binding.

To assign the detected cross-links to a structural model, we analysed the satisfaction rate of the SDA Mic60-Mic19 dataset with the closed and open putative conformations, revealing that the complex dataset correlates better with the open, hetero-octameric Mic60-Mic19 model (Supplementary Figure 8A, C). Mapping the Mic60-Mic19 SDA cross-linking data onto the two models confirms that the majority of the intramolecular mitofilin-CC contacts seen in the *apo* dataset were lost, supporting the idea that the extended mitofilin region dissociates from the CC and adopts an open conformation upon Mic19 binding (Figure 6B, Supplementary Figure 8A). This rearrangement enables the IH of Mic19 to occupy the Mic60 CC tail previously engaged by the EHR, as confirmed by multiple SDA cross-links between them, when Mic19 was incorporated into the ‘open’ model (Figure 6B right, box 3). Interestingly, the Mic60-Mic19 complex dataset also contains cross-links between mitofilin domains that are compatible with the domain-swapped dimeric form, supporting a model in which both monomeric and dimeric conformations coexist (Supplementary Figure 8B). In summary, the *in vitro* cross-linking data support a model in which Mic60 transitions from a closed, *apo* dimer to an open, Mic19-bound hetero-octamer, with the EHR acting as the central structural switch of this conformational change (Figure 6C).

### Mic60 and Mic19 undergo large conformational rearrangements upon complexation

The cross-linking data reveal a pronounced structural rearrangement within Mic60 upon Mic19 binding, consistent with a reorganization of intramolecular contacts accompanying complex assembly. To further address such a potential mechanism, we aimed to obtain molecular insights into the conformational dynamics of Mic60-Mic19 complex formation. To this end, we performed hydrogen-deuterium exchange mass spectrometry (HDX-MS) for Mic60_sol_ and for full-length Mic19, both in isolation and when present in a complex. HDX differences in the isolated and assembled proteins were then plotted on the detected peptides of the proteins.

In these experiments, prevalent changes in deuterium exchange were detected in several regions of both proteins upon complex formation (Figure 7A, B, black trace). As expected, the experimentally described CHCH-mitofilin domain (Site 1a) interface showed a marked decrease in deuterium incorporation upon complex formation (Figure 7A, C). HDX coverage for the CHCH domain in Mic19 was low. This may be explained by its stable disulfide bond, which is known to cause reduced detection of peptides in their vicinity (Mysling et al., 2014). The predicted IH-CC tail interface (Site 2) was also shielded from solvent exchange upon complex formation, with marked changes in the percentage of deuterium incorporation in Mic60 CC and Mic19 IH (Supplementary Figure 9C).

**Figure 7:**
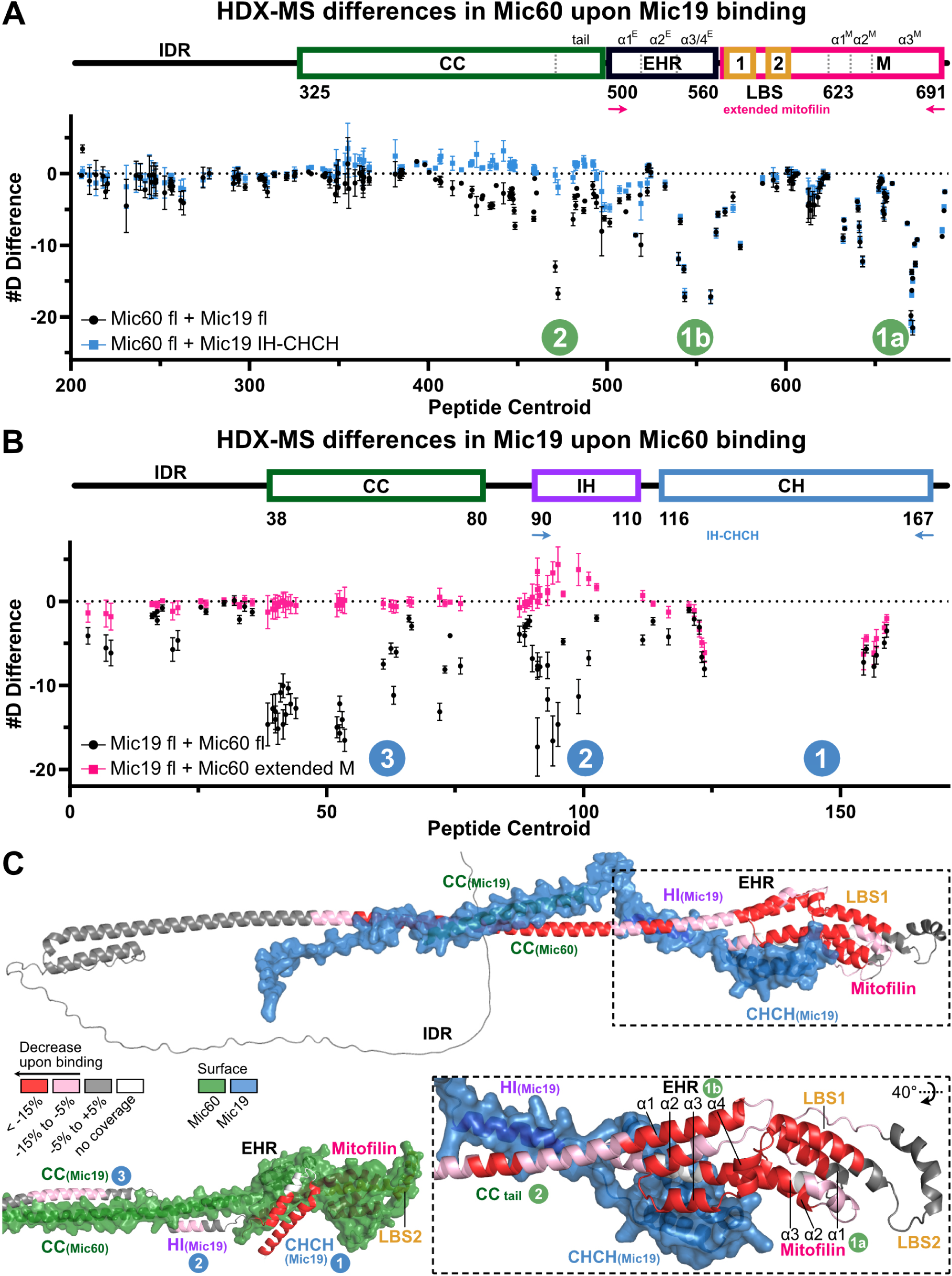
HDX-MS reveals extensive conformational rearrangements across the Mic60 sequence upon Mic60-Mic19 complexation. **A** Differential deuterium exchange in Mic60 upon Mic19 binding, either with full-length Mic19 (black circle) or with a truncated construct IH-CHCH (blue square), and **B** in Mic19 upon Mic60 binding, either with full-length Mic60 (black circle) or with the extended mitofilin domain (pink square). Each point represents a single peptide, mapped on the peptide centre, with the standard deviation of triplicates marked as bars. A graphic illustration as in Figure 4A annotates the domains corresponding to the residues on the x axis. The regions showing large differences in deuterium incorporation are numbered in green for Mic60 and in blue for Mic19, corresponding to Sites 1 and 2 from Figure 4A and an additional site 3 for Mic19. **C** Peptides that showed significant differences in HDX as shown in Supplementary Figure 9A, B were mapped onto the AlphaFold2 prediction from Figure 4A, with the interacting regions marked by circles as in A and B. When Mic60 shifts upon Mic19 binding are plotted (top and bottom right), Mic19 is shown as a blue surface. When Mic19 HDX shifts are plotted (bottom left), Mic60 is shown as a green surface.

Of note, the EHR appeared to be stratified into two submodules with distinct local binding dynamics (Figure 7A). Helices α1^E^ and α2^E^ displayed only a limited decrease in deuterium exchange upon complex formation (Site 1b), consistent with their solvent protection through tight packing as modelled for both the *apo* and Mic19-bound states (Figure 6B). In contrast, helices α3^E^ and α4^E^ (Site 1b) exhibited near-complete deuterium incorporation in the *apo* state (Supplementary Figure 9C) but were shielded in the Mic19-bound state, similar to the CHCH-mitofilin and the IH-CC tail interactions. These data therefore support the notion of the dynamic nature of the EHR, undergoing extensive packing rearrangements within Mic60 upon Mic19 binding. Helices α3^E^-α4^E^ may stabilize the EHR in its Mic19-associated state through residues V546/A549, whereas helices α1^E^-α2^E^ may primarily facilitate conformational switching, which ultimately allows for the essential secondary binding site between the CC tail and the IH to form.

Interestingly, HDX-MS analysis of truncation constructs containing only the IH-CHCH (Figure 7A, blue trace) or the extended mitofilin region (Figure 7B, pink trace), reveals the same protection pattern at the CHCH-mitofilin interface of site 1, but not in the IH-CC tail binding in site 2. This supports the idea that Mic60 aligns with Mic19 in a parallel, zipper-like fashion from the C- to the N-terminal end, and that complex formation is initiated by CHCH-mitofilin association, with the IH-CC tail forming a secondary binding site in the full-length proteins. In the absence of this binding site, the Mic60 subcomplex cannot assemble fully to its hetero-octameric form, as shown by BN-PAGE (Supplementary Figure 9D).

Taken together, the HDX-MS findings correlate both with the cross-linking MS analysis and the structural models presented here, describing Mic60 subcomplex assembly as a highly dynamic and distributed process, spanning the CHCH-mitofilin and IH-CC tail interfaces.

## Discussion

Here, we report that Mic60, the core subunit of MICOS, undergoes a drastic conformational rearrangement during complex formation with Mic19 to govern formation of mitochondrial crista junctions. The structurally defined EHR of the extended Mic60 mitofilin domain acts as a molecular switch driving complex formation with Mic19 through a distinct interaction site. A further binding site between the two proteins was identified between the Mic19 IH and the C-terminal end of the coiled-coil (CC tail). By an integrated approach combining structural, biochemical, biophysical and cell-based data, we characterize the dynamic assembly of the Mic60 subcomplex and the role of the EHR as a determinant for mitochondrial crista junction formation and therefore mitochondrial architecture and homeostasis.

To study the mechanistic basis of Mic60-Mic19 complex formation, we determined the crystal structure of an extended mitofilin construct containing an N-terminal extension, the EHR. In this structure, the EHR closely associates with the core mitofilin domain and contributes to Mic19 binding, consistent with an integral and conserved role in the Mic60-Mic19 assembly mechanism. This structural view is supported by a recent evolutionary analysis of ancestrally reconstructed Mic60 orthologues (Benning et al., 2025) and our own studies (Supplementary Figure 2), which identify the EHR as an evolutionarily conserved module of an ancient extended mitofilin domain.

While our structure confirms the previously published Mic60-Mic19 interaction interface involving the mitofilin and CHCH domains, it reveals a 1:1 hetero-dimer rather than the previously reported domain-swapped 2:2 hetero-tetramer (Bock-Bierbaum et al., 2022). In the absence of a full-length Mic60 structure, we cannot unambiguously distinguish between the monomeric and dimeric assembly of the Mic60 mitofilin domain in the apo and Mic19-bound states. However, cross-linking data indicate close proximity of two mitofilin domains in both states. Notably, both available structures of the mitofilin domain are derived from truncation constructs that retain Mic19-binding activity but are insufficient to support assembly of the complete hetero-octameric complex, as shown in BN-PAGE experiments (Supplementary Figure 9D). Therefore, full Mic60 subcomplex assembly requires interactions beyond the minimal Mic60-Mic19 binding interface.

Seeking to characterize these additional interactions, we identified the EHR as critical determinant for Mic19 binding. In particular, two highly conserved EHR residues (V546 and A549 in the *Chaetomium* sequence) directly engage in Mic19 binding *in vitro* and *in vivo*, arguing for an evolutionary conserved function. We also identified a further interaction site between the two proteins involved in the Mic19 IH and the Mic60 coiled-coil. Although this interface is required for Mic60-Mic19 assembly *in vitro*, *in vivo* experiments indicate a more subtle role for Mic60 subcomplex assembly compared to the mitofilin-CHCH interface.

While the EHR extensively interacted with the core mitofilin domain in the current structure, it bound to the Mic60 coiled-coil tail region in a previous crystal structure (Bock-Bierbaum et al., 2022) through the same interface. Cross-linking and HDX data presented here suggest that the EHR associates with the CC in the absence of Mic19, although the atomic structure of this state remains unresolved. We propose that this interaction brings the mitofilin domain into close proximity to the CC and leads Mic60 to adopt a closed, dimeric conformation in the absence of Mic19, covering the tetramerization interface and therefore blocking Mic60 tetramer assembly. With the mitofilin domain being disengaged from the EHR, the closed Mic60 architecture may accommodate a domain-swapped mitofilin dimer, which is spatially compatible with our dimeric model of the apo Mic60 state (Supplementary Figure 7B).

Our findings identify the EHR as a dynamic module that switches between the Mic19-free inactive state and the Mic19-bound active state. Consistent with this model, we observed striking differences in the cross-linking pattern of full-length Mic60 before and after Mic19 binding, centering around the EHR as a critical hinge. In the absence of Mic19, the extended mitofilin region cross-linked extensively with the CC, consistent with a closed architecture. Upon Mic19 binding, these cross-links were largely absent, supporting a model in which the extended mitofilin disengages from the CC and Mic60 adopts a more open conformation. The open model was further corroborated by several cross-links between various domains of Mic60-Mic19. Furthermore, HDX-MS measurements recapitulate both the mitofilin-CHCH and IH-CC tail binding sites, reflecting these vast structural rearrangements and illustrating the dynamic nature of Mic60-Mic19 binding.

Our model of the EHR structural switch could rationalize the regulation of Mic60-Mic19 subcomplex assembly through post-translational modifications. In *Drosophila melanogaster*, phosphorylation by the mitochondrial kinase PINK1 is thought to be critical for Mic60 stability at CJs, and the corresponding phosphorylation site aligns with residue S520 in the EHR (Supplementary Figure 2A) (Tsai et al., 2018), along with other well-conserved EHR residues that have been identified in multiple phosphorylation screens (Hornbeck et al., 2015). Modification of these residues could influence Mic19 binding. Since they are placed close to the positively charged EHR-LBS1 interface (Supplementary Figure 3C), they could also weaken membrane binding. In this way, complex assembly and CJ formation could be dynamically regulated in response to external stimuli.

Integrating our findings with existing structural data, we propose an activation mechanism for Mic60 in which the EHR functions as a conformational switch that drives Mic60 from a closed, dimeric form to its hetero-octameric assembly with Mic19 (Figure 8, Supplementary Video 1). In the closed state, the EHR packs against the CC tail, positioning the mitofilin across the CC in a compact protein conformation and thereby blocking tetramerization. Mic19 binding then triggers a zipper-like assembly mechanism, initiated by the high-affinity mitofilin-CHCH interaction, and then further involving the EHR-CHCH and the IH-CC tail interfaces. EHR release from the CC leads to an opening of Mic60 and the CC tetramerization interface becomes accessible, allowing the formation of the hetero-octameric Mic60-Mic19 complex at the CJ (Supplementary Video 1). Finally, binding of the Mic19 N-terminus to the OMM-localized SAM complex (Tang et al., 2020) could anchor the Mic60-Mic19 complex to the OMM.

**Figure 8:**
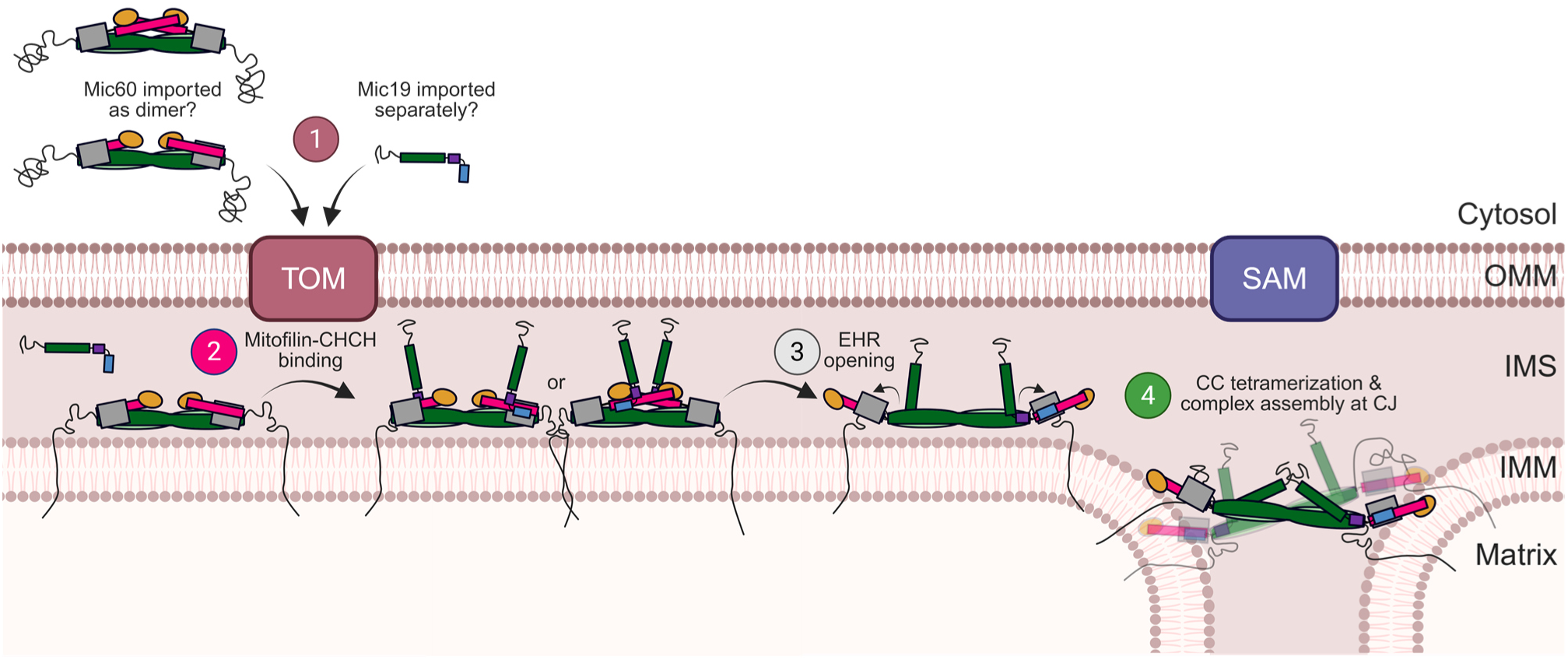
Proposed assembly model of the Mic60-Mic19 complex. Schematic overview of the hypothesized assembly of the Mic60-Mic19 complex in CJs. (1) Mic60 and Mic19 are imported into the mitochondrion via the translocase of the outer membrane (TOM) pathway, possibly separately, and Mic60 is inserted into the IMM. In the closed, autoinhibited dimer, the CC tail is occupied by the EHR. (2) In the presence of Mic19, the CHCH domain binds to the mitofilin domain, in a monomeric or dimeric form. (3) This drives a switch motion which releases the EHR from the CC tail, with the EHR now associating with the core mitofilin domain to form the extended mitofilin region. This open conformation allows the IH of Mic19 to bind the previously occupied Mic60 CC tail. (4) This conformation further frees the tetramerization interface of the Mic60 CC, allowing another Mic60-Mic19 hetero-tetramer to associate (here illustrated transparently), and assembling the hetero-octameric Mic60-Mic19 complex at the CJ. The Mic19 N-terminus could possibly arch over the IMS and contact SAM (distances not drawn to scale). Created in BioRender.

Our mechanistic framework for Mic60 subcomplex assembly has direct implications for CJ biogenesis. Once assembled, the Mic60 subcomplex may stabilize newly formed CJs by cross-linking and bridging opposing IMM sheets (Harner et al., 2016; Stephan et al., 2020). In parallel, assembly of the Mic10 subcomplex into ring-like structures at the CJ neck may contribute to the formation of ring-like CJs (Brown et al., 2026). In this way, the Mic60 activation mechanism described here provides a molecular link between the dynamic assembly of MICOS subcomplexes and CJ formation, offering a mechanistic framework for how MICOS establishes and maintains mitochondrial membrane architecture.

## Materials and Methods

### Cloning and plasmids

*ct*Mic60 (UniProt ID: G0SHY5) and *ct*Mic19 (UniProt ID: G0S140) were used for cloning via restriction sites *Nde*I or *BamH*I and *Xho*I and subsequent ligation into a modified pET28a vector encoding an N-terminal His_6_ tag fusion protein as described previously (Bock-Bierbaum et al., 2022; Hessenberger et al., 2017). Variants were introduced via site-directed mutagenesis (Ho et al., 1989).

### Expression and purification

Expression plasmids were freshly transformed into BL21(DE3) *Escherichia coli* cells, and transformed bacteria were grown in terrific broth media supplemented with 100 μg/ml kanamycin at 37°C and 80 rpm until they reached an OD_600_ of 0.8. Protein expression was induced with 300 μM isopropyl-ß-D-1-thiogalactopyranoside (IPTG, Carl Roth) and the cells were cooled to 18°C and incubated overnight. Cells were harvested via centrifugation for 20 min at 4,000 x *g* and cell pellets were stored at −20°C.

Cell pellets were thawed and resuspended in lysis buffer (20 mM HEPES/NaOH pH 7.5, 500 mM NaCl, 20 mM imidazole), supplemented with 500 μM 4-(2-aminoethyl)-benzolsulfonyfluorid hydrochloride (AEBSF, BioChemica) and 1 μM DNAse (Roche). Cells were lysed through sonication (5x 45 s with 1 second pulses at 70% power) and lysates were centrifuged at 40,000 x *g* for 45 min in an Avanti J-26 XP centrifuge using a JA-25.50 rotor (Beckman). The supernatant was filtered using a 0.45 μM filter and applied to Ni Sepharose High Performance resin (GE Healthcare), equilibrated in lysis buffer. The column was washed with 5 column volumes (CV) lysis buffer and 5 CV wash buffer (20 mM HEPES/NaOH pH 7.5, 500 mM NaCl, 40 mM imidazole) and the protein was eluted in 5 CV elution buffer (20 mM HEPES/NaOH pH 7.5, 500 mM NaCl, 500 mM imidazole). Glutathione-S-Transferase (GST)-His_6_-tagged human rhinovirus 3-chymotrypsin protease (HRV 3C) was added at a 1:50 ratio (w/w) and the protein mixtures were dialyzed overnight at 4°C to low imidazole buffer (20 mM HEPES/NaOH pH 7.5, 150 mM NaCl, 20 mM imidazole). The protein was subsequently applied to a reequilibrated Ni-NTA column to remove the residual His_6_-tag and 3C protease. The flow-through was collected, concentrated using an Amicon filter concentrator of 3 kDa, 10 kDa or 30 kDa cut-off (Millipore) and applied on a HiLoad™ 16/600 Superdex™ 75 or 200 or Superose™ 6 preparative grade column connected to an ÄKTA Pure chromatography system (Cytiva). Protein was eluted with Size Exclusion Chromatography buffer (SEC, 20 mM HEPES/NaOH pH 7.5, 150 mM NaCl) at a flow-rate of 1 ml/min and 1 ml fractions were collected. Pure protein fractions were pooled, concentrated, flash-frozen in liquid nitrogen and stored at −70°C. For deletion variants or point variants of the Mic60 extended mitofilin domain the N-terminal His_6_ tag was not removed during purification.

The construct extended mitofilin was purified using an on-column cleavage protocol. Lysis was performed as described and the supernatant was applied to Ni Sepharose High Performance resin (GE Healthcare), equilibrated in lysis buffer. The column was washed with 5 CV lysis buffer before the beads were resuspended in 2 CV lysis buffer and 0.1 mg of HRV 3C protease was added to the sample and incubated under gentle rotation at 4°C overnight. The following day the sample and beads were applied to the column again. The flow-through was collected and the Ni-NTA beads were washed first with 5 CV of lysis buffer and secondly with 5 CV of elution buffer to remove the residual protein tag. The SEC was performed as described above.

For all protein samples, purity was confirmed by mass spectrometry as described previously (Bock-Bierbaum et al., 2022) and samples were flash-frozen in liquid nitrogen and stored at −70°C.

### Crystallization, data collection, refinement and structural analysis tools

The constructs extended mitofilin and IH-CHCH were expressed and purified separately. 2.2 mg of each protein were mixed, diluted to 500 μl using SEC buffer, incubated on ice for 30 min and subsequently co-eluted using a HiLoad™ 16/600 Superdex™ 200 preparative grade column connected to an ÄKTA Pure chromatography system (Cytiva), using a flow rate of 1 ml/min and SEC buffer composed as above. Crystallization trials were performed using the sitting-drop vapor-diffusion method in a 96-well crystallization plate using a Gryphon pipetting robot (Matrix Technologies Co.) by mixing 200 nl protein with 200 nl reservoir solution and equilibrating against 80 μl reservoir. Commercial screens, namely JBScreen Basic HTS, JBScreen JCSG+ HTS (Jena Bioscience), Classics II Suite, pH Clear Suite I and II and Protein Complex Suite (Qiagen), were set up at 4°C and 20°C using a concentration of 10.9 mg/ml. The Rock Imager 1000 storage system (Formulatrix) was used for storing and imaging the experiments.

Initial hits in the form of a crystal shower appeared on day 7 under the conditions 0.2 M Li_2_SO_4_, 0.1 M Tris/HCl pH 8.5, 30% (w/v) PEG 4000 at 20°C. For all further optimization steps the protein concentration was increased to 15.4 mg/ml. Optimizing the PEG concentration to 25-27.5% while retaining the salt and pH led to individual crystal cluster formation, but the crystals were amorphous, non-uniform and small in size (maximally 50 μm). To further improve crystal morphology and acquire larger singular crystals rather than a crystal shower, microseeding was performed. Briefly, a seed bead was incubated with crystallization cocktail (0.2 M Li_2_SO_4_, 0.1 M Tris/HCl pH 8.5, 27.9% (w/v) PEG 4000) and purified protein. Crystallization drops containing crystal hits, namely needles and clusters, were added to the bead and were vortexed for 1 min to crush the crystals into microcrystalline. Two crystallization plates were dispensed: the optimization screen described above, which gave rise to the amorphous clusters, and Classics Suite II. On day 49 crystal growth was observed in one of the conditions of Classics Suite II (0.2 M Li_2_SO_4_, 0.1 M HEPES/NaOH pH 7.5, 25% (w/v) PEG 3350), where multiple crystal needles were forming a bundle from one nucleation centre. The needles continued growing in length until day 108, at which point they had reached a length of 600 μm and were fished using 20% ethylene glycol as cryo-protectant.

Diffraction data were collected at a wavelength of 0.9184 Å and −173°C on beamline 14.1 operated by the Helmholtz-Zentrum Berlin at the BESSY II synchrotron in Berlin Adlershof (Mueller et al., 2025; Mueller et al., 2015). 2,000 images were collected using 0.25 s exposure time at a detector distance of 425.62 mm and an oscillation increment of 0.1°. Data was automatically indexed, integrated and scaled using XDSAPP V. 3.1.9c (Sparta et al., 2016). The extended M IH-CHCH complex crystallized in space group P3_1_21 (152).

The crystallized constructs showed high sequence coverage (62%) by the previously solved dimeric structure of the *ct*Mic60 LBS1-mitofilin and *ct*Mic19 CHCH domain (PDB ID: 7PV1). However, efforts to solve the phase problem through MR with this structure, either as a monomer or as a dimer, failed. AlphaFold was used to predict the novel Mic60-Mic19 complex structure, which was predicted with high confidence except for the Mic19 N-terminal residues 90-114 (Jumper et al., 2021). The residues were deleted and the crystallographic phase problem was solved by molecular replacement using this modified prediction of the hetero-dimer with Phaser-MR implemented in the Phenix distribution (V. 1.20.1_4487) (Liebschner et al., 2019; McCoy et al., 2007). The structure was solved using iterative cycles of Refmac5 in the CCP4 suite followed by Coot V. 1.1.11 for manual model building (Emsley et al., 2010; Murshudov et al., 1997). A crucial step was the deletion of *ct*Mic60 residues 500-545 in the beginning of refinement and their later placement as polyalanine helices, which were eventually mutated to the protein sequence. Once all visible residue backbone, sidechains and waters had been built the model was refined using phenix.refine, applying individual B factors, secondary structure restraints and Translation-Liberation-Screw rotation (Afonine et al., 2012). The final model appeared as a hetero-dimer of *ct*Mic60 aa 504-586-GS-622-691 bound to *ct*Mic19 aa 117-164. The extended mitofilin domain construct had a very similar structure to the AlphaFold2 prediction (Supplementary Figure 1E). In Mic19, the IH was not visible, pointing to a high flexibility of this region or the possibility it was degraded prior to crystal formation. N-terminally extended constructs of Mic60, *apo* or bound to Mic19, failed to yield crystals. Final model validation and data statistics were performed with Molprobity V. 4.0.2 (Williams et al., 2018) and can be seen in Table 1. The structure was deposited and validated in the PDB database under PDB ID 28XR.

Figures were generated using PyMol (V. 2.5.5) (Schrödinger & DeLano 2020), while cross-links were visualized on models using ChimeraX with the XMAS tool (Lagerwaard et al., 2022; Meng et al., 2023). PDBePISA was used for interface area calculation of the *ct*Mic60-*ct*Mic19 hetero-dimer (Krissinel & Henrick, 2007). Electrostatics were calculated on the Poisson-Boltzmann server, using PDB2PQR followed by APBS (Jurrus et al., 2018). Conservation plots were generated using ConSurf Web server using default parameters (Yariv et al., 2023). Conservation of Mic19 was calculated based on the full-length sequence, while for Mic60 only the residues of the extended mitofilin region were provided as input, as slight differences in the conservation of the Mic60 C-terminus were obscured when calculated for the entire *ct*Mic60 sequence due to the overall high conservation of the C-terminus (see Supplementary Figure 2). The aligned Mic60 sequence segments were extracted and imported to Jalview (Waterhouse et al., 2009) to generate a conservation score plot.

### Isothermal titration calorimetry (ITC)

ITC measurements were performed using a MicroCal PEAQ-ITC instrument (Malvern) according to the manufacturer’s instructions at 25°C. The injection programme of 12 injections of 3 μl was used. Samples were dialysed overnight into SEC buffer prior to measurement. *ct*Mic19 constructs were placed in the syringe (450 μM, except for *ct*Mic19 I102D at 400 μM and *ct*Mic19 90-164 IH-CHCH at 300 μM) and *ct*Mic60 constructs were placed in the cell (45 μM). Results were analysed using the MicroCal PEAQ-ITC Analysis Software (Malvern). As in previous experiments (Bock-Bierbaum et al., 2022), the binding numbers were often lower than 1, which may be related to the heterogeneous oligomeric states of Mic60 that may not always yield a 1:1 binding or some Mic60 precipitation during the ITC measurements.

### Analytical size-exclusion chromatography (SEC) coupled to right-angle light scattering (SEC-RALS)

Analytical SEC-RALS was performed using an Agilent HPLC system equipped with an autosampler G1329B and an Omnisec Reveal (Malvern), connected to a Superdex™ 200 Increase 5/150 GL column (Cytiva). The column and RALS instrument were equilibrated overnight with SEC buffer (20 mM HEPES/NaOH pH 7.5, 150 mM NaCl). 100 μl sample at a concentration of 2 mg/ml (incubated on ice for 30 min in the case of extended mitofilin_IH-CHCH complexation) was injected using the autosampler and eluted at a flow rate of 0.2 ml/min at 20°C. UV absorption at 280 nm, 260 nm, 210 nm and light scattering were measured and analysed by the Agilent Offline and Malvern Omnisec software.

### Analytical size-exclusion chromatography (SEC) for Mic60-Mic19 complex formation

Analytical SEC was performed using an Agilent HPLC system equipped with an autosampler G1329B, connected to a Superose™ 6 Increase 5/150 GL column (Cytiva). The column was equilibrated overnight with SEC buffer (20 mM HEPES/NaOH pH 7.5, 150 mM NaCl). 100 μl sample (*ct*Mic60_sol_ variants with a concentration of 1 mg/ml, *ct*Mic19 variants with a concentration of 0.36 mg/ml, incubated on ice for 30 min) was injected using the autosampler and eluted at a flow rate of 0.2 ml/min at 20°C. UV absorption at 280 nm, 260 nm, 210 nm were measured and analysed by the Agilent Offline software.

### Blue Native polyacrylamide gel electrophoresis (BN-PAGE)

Recombinant proteins were analysed by BN-PAGE as described previously (Bock-Bierbaum et al., 2022). Briefly, 5 μg of *ct*Mic60 and an equimolar amount of *ct*Mic19 (1.8 μg) were incubated on ice for 30 min. All samples were diluted to 10 μl using SEC buffer (20 mM HEPES/NaOH pH 7.5, 150 mM NaCl), 2.5 μl NativePAGE™ Sample Buffer (Invitrogen) was added and the total sample volume was loaded on a 4-16% acrylamide gradient BN-PAGE gel (Invitrogen). 5 μl NativeMark™ Unstained Protein Standard (Invitrogen) were loaded as reference and the gel was run at 150 V for 2 h on ice and subsequently stained using Coomassie Brilliant Blue.

### *S. cerevisiae* strains and plasmids

All *Saccharomyces cerevisiae* strains used in this study were derivatives of YPH499 and were based on either the WT strain, Mic60-Protein A *Δmic19*, or Mic19-Protein A *Δmic60* strains, which have been described previously (Bohnert et al., 2015; von der Malsburg et al., 2011). Plasmids pRS416 and pRS426 harboring the open reading frames of either *MIC60* or *MIC19*, including their native promoters and terminators, have also been described previously (Bock-Bierbaum et al., 2022; Bohnert et al., 2015). Point mutations in the respective genes were introduced by site-directed mutagenesis. Wild-type and deletion strains were transformed with either the corresponding empty vector or plasmids expressing *MIC60* or *MIC19* variants. Yeast cells were grown in liquid minimal glycerol medium (0.67% (w/v) yeast nitrogen base, 0.07% (w/v) complete supplement mixture (CSM) lacking uracil, and 3% (v/v) glycerol) at 30°C.

### Isolation of *S. cerevisiae* mitochondria and IgG affinity purification

Cells were grown in liquid minimal glycerol medium to mid-logarithmic phase and harvested by centrifugation at 1,200 g for 5 min at RT. Cell pellets were resuspended in DTT softening buffer (0.1 M Tris/H_2_SO_4_ pH 9.4, 10 mM DTT) at a ratio of 2 ml per gram wet cell weight and incubated for 20 min at 30°C. Following centrifugation at 2,000 x *g* for 5 min at RT, cells were washed with Zymolyase buffer (1.2 M sorbitol, 20 mM KPi pH 7.4) and subsequently resuspended in the same buffer containing 4 mg Zymolyase per gram wet cell weight. After incubation for 30 min at 30°C to generate spheroplasts, cells were collected by centrifugation at 2,000 g for 5 min at RT, washed once with Zymolyase buffer, and resuspended in homogenization buffer (0.6 M sorbitol, 10 mM Tris/HCl pH 7.4, 1 mM EDTA, 0.2% (w/v) bovine serum albumin, 1 mM phenylmethylsulfonyl fluoride (PMSF)) at a ratio of 6.5 ml per gram wet cell weight. Spheroplasts were disrupted using a glass-Teflon Dounce homogenizer (15 strokes). Cell debris was removed by centrifugation at 1,500 g for 5 min at 4°C. The resulting supernatant was subjected to a second centrifugation step at 15,000 x *g* for 10 min at 4°C to pellet mitochondria. The mitochondrial pellet was resuspended in SEM buffer (250 mM sucrose, 10 mM MOPS pH 7.2, 1 mM EDTA). Protein concentrations were determined using the Bradford assay and adjusted to 10 mg total protein/ml. Isolated mitochondria were flash-frozen in liquid nitrogen and stored at −80°C until use.

For IgG affinity purification, 0.9 mg of isolated *S. cerevisiae* mitochondria harboring Mic60-Protein A or Mic19-Protein A fusion proteins were solubilized in solubilization buffer (20 mM Tris/HCl pH 7.4, 50 mM NaCl, 0.1 mM EDTA, 10% (v/v) glycerol, 2 mM PMSF, 1% (w/v) digitonin) for 30 min at 4°C. Insoluble material was removed by centrifugation at 20,000 x *g* for 10 min at 4°C. The resulting mitochondrial extracts were incubated with 50 μl human IgG-Sepharose beads, pre-equilibrated with 0.5 M acetate (pH 3.4), for 90 min at 4°C. Following binding, beads were washed ten times with digitonin buffer (20 mM Tris/HCl pH 7.4, 0.5 mM EDTA, 60 mM NaCl, 10% (v/v) glycerol, 2 mM PMSF, 0.3% (w/v) digitonin), with centrifugation at 700 x *g* for 30 s at 4°C between washing steps. Bound proteins were eluted by overnight cleavage with tobacco etch virus (TEV) protease at 4°C, followed by centrifugation at 1,200 x *g* for 30 s. Eluates were mixed with trichloroacetic acid for protein precipitation and, together with the samples of the original mitochondrial extract (totals), analysed by SDS-PAGE and western blotting.

### Electron microscopy of *S. cerevisiae* mitochondria

*S. cerevisiae* cells grown in liquid minimal glycerol medium (0.67% (w/v) yeast nitrogen base, 0.07% (w/v) complete supplement mixture (CSM) lacking uracil, supplemented with 0.1% (w/v) glucose and 3% (v/v) glycerol) for 24 hours at 30°C until early log phase and handled as previously described (Bock-Bierbaum et al., 2022). Cells were fixed using 4% (w/v) paraformaldehyde and 0.5% (v/v) glutaraldehyde (GA) in 0.1 M citrate buffer, freshly prepared, for 3 hours at 30°C. They were washed with citrate buffer and permeabilized with 1% (w/v) sodium metaperiodate for 1 hour at 4°C. Cells were washed with 0.1 M phosphate buffer before being embedded in 12% gelatine: after 10 min of incubation, the 37°C warm gelatine was cooled on ice. 1 mm^3^ cubes were cut and infiltrated overnight with 2.3 M sucrose, and they were mounted onto specimen chucks and flash-frozen in liquid nitrogen. Ultrathin sections were cut using an UC7 ultramicrotome (Leica) at −110°C and collected on formvar/carbon-coated copper grids (Plano). The gelatine was removed before staining with 3% (w/v) silicotungstic acid hydrate (Fluka) in 2.8% (w/v) polyvinyl alcohol (Sigma-Aldrich) in water for 5 min. Grids were imaged with the transmission electron microscope JEM 2100 Plus (Jeol) operating at 200 kV and equipped with a 20MP CMOS Xarosa camera and the imaging software Radius V2.2 (Emsis) at ×20,000 magnification. Data were collected, shuffled and anonymised using the Microscopy Image Browser (Belevich et al., 2016) and were evaluated by three independent individuals. All data are presented as the average value, rounded to a whole number, with means ± standard deviation of 100 mitochondria per condition plotted, and value differences were compared statistically. Data analysis and plotting were done with the statistic program GraphPad Prism V9 (Dotmatics). The normal distribution was tested using the Shapiro-Wilk test and the data were not normally distributed. Since the data were also unpaired, the two-sample nonparametric Kolmogorov-Smirnov was used. For all groups, *n* = 100 mitochondrial cross sections were analysed. Differences of *p* ≤ 0.05 were considered significant (*p < 0.05, \*\**p* < 0.01, and ***p < 0.001).

### *In vitro* cross-linking coupled to LC-MS

50 μg *ct*Mic60 (*apo)* as well as 50 μg *ct*Mic60 in complex with an equimolar amount of *ct*Mic19 (17.9 μg, complex) were cross-linked with 0.5, 1 or 2 mM Succinimidyl 4,4’-azipentanoate (SDA, Sigma-Aldrich) at a protein concentration of 1 mg/ml and incubated for 45 min at RT, shaking. Samples were then irradiated for 30 s at 365 nm with 100% intensity (LED Cube 100 IC, Dr. Hönle AG UV Technology) and subjected to in-solution trypsin digestion.

Cross-linked samples were treated with 8 M urea, 5 mM Tris(2-carboxyethyl)phosphine hydrochloride (TCEP), and 40 mM chloroacetamide and the mixture was incubated at 37 °C for 1-1.5 hours in the dark. Proteolytic digestion was performed using LysC (1:50 ratio (w/w)) and trypsin (1:50 ratio (w/w)). The solution was then diluted to a final concentration of 2 M urea using 50 mM triethylammonium bicarbonate buffer (pH 8.5). The digestion was stopped with 1% formic acid, and peptides were desalted using C8 StageTips and dried before being stored at −20°C until measurement.

Peptides were resuspended in 1% acetonitrile with 0.05% trifluoroacetic acid and 1 µg was injected into the same LC-MS system used for disulfide mapping. Detailed acquisition parameters are listed in Supplementary Table 2.

RAW files were identified using XlinkX (Liu et al., 2017), implemented in Proteome Discoverer versions 3.2, operating in open-search mode. Searches were conducted against a database containing the target protein sequences. The false discovery rate was controlled at 1%, with separate estimation for intra- and inter-links. Dynamic score thresholds were applied based on decoy CSM scores, and identifications were further filtered post-search by minimum CSM count. All cross-links were visualized using xiNET (Combe et al., 2015) and ChimeraX with the XMAS tool (Lagerwaard et al., 2022; Meng et al., 2023).

### Assembly of the *ct*Mic60 dimer and *ct*Mic60-*ct*Mic19 hetero-octamer models

To visualize the conformational rearrangements suggested by the cross-linking data, we assembled two structural models of *ct*Mic60. Both models were constructed based on two available structures of fungal Mic60: the CC domain of *Lachancea thermotolerans* Mic60 (PDB ID: 7PUZ, residues 207-382, equivalent residues in *ct*Mic60 are residues 402-539) and the Mic60 extended mitofilin domain structure bound to the Mic19 CHCH domain described here (*ct*Mic60 aa 504-586-GS-622-691 bound to *ct*Mic19 aa 117-164). A homologous model of the CC domain was generated using SWISS-MODEL, while preserving its tetrameric assembly (Waterhouse et al., 2018). To incorporate the flexible LBS2 (*ct*Mic60 aa 587-621) we extracted it from the AlphaFold2 full-length *ct*Mic60-*ct*Mic19 prediction described above (Jumper et al., 2021) and manually fused it to the extended mitofilin structure using Coot V. 1.1.11 (Emsley et al., 2010). For Mic19, we extracted the IH-CHCH segment of the same AlphaFold2 prediction (*ct*Mic19 aa 90-164), excluding the IDR and CC as structural informational on their conformational dynamics remains limited.

All models comprise *ct*Mic60 residues 402-691 and, where applicable, *ct*Mic19 residues 90-164. The key difference between them lies in the overall conformation of Mic60 and more specifically of the EHR. In the “closed” model, we sought to represent the Mic60 *apo* state, and therefore excluded Mic19. Here, EHR helices α1^E^ and α2^E^ (*ct*Mic60 aa 504-539) adopt a conformation analogous to that observed in the ltMic60 CC structure, forming a four-helix bundle with the CC termini (equivalent to *ct*Mic60 aa 410-430 and 480-500). The extended mitofilin domain is positioned in close proximity to the CC, giving rise to the designation “closed conformation”. As this is not an experimentally solved structure, the exact positioning of the remaining EHR helices α3^E^ and α4^E^ and the mitofilin domain is not resolved in detail and is rather intended to represent the overall architecture of the C-terminus. If the extended mitofilin region forms a monomer as in 28XR and Supplementary Figure 7A, this results in clashes between the central mitofilin helix α3^M^ and the CC. Alternatively, the EHR helices can remain associated with the CC tail, while the mitofilin domain forms the domain-swapped dimer, as in 7PV1 and Supplementary Figure 7B.

In the second model, we aimed to represent the Mic19-bound state. In this “open” conformation, the EHR adopts the conformation observed in the extended mitofilin region structure, bound to the CHCH domain of Mic19. Here, the EHR dissociates from the CC tail, and the same vacated region is now instead occupied by the Mic19 IH. This rearrangement repositions the extended mitofilin region away from the CC, thereby relieving steric occlusion. As a consequence of this opening, the tetramerization interface of the CC becomes accessible, allowing the incorporation of two additional copies of Mic60 and Mic19 to the complex and yielding a hetero-octameric assembly for the model.

### Hydrogen-Deuterium Exchange Mass Spectrometry (HDX-MS) measurements

*ct*Mic60 and *ct*Mic19 constructs were eluted on a HiLoad™ 16/600 Superdex™ 75 or 200 or Superose™ 6 preparative grade column connected to an ÄKTA Pure chromatography system (Cytiva) either in the *apo* state or as co-eluted complexes in SEC buffer (20 mM HEPES/NaOH pH 7.5, 150 mM NaCl) and analysed using SDS-PAGE.

Deuterium exchange reactions were induced upon the addition of 45 μl deuterium buffer (20 mM HEPES/NaOH pH 7.5, 150 mM NaCl, 93% D_2_O) to 5 μl of pre-diluted protein at 5 μM, leading to a final concentration of 0.5 μM protein. The reactions were quenched by the addition of 20 μl ice-cold quench buffer (final concentration: 0.57 M guanidinium chloride, 0.86% formic acid). Samples were snap-frozen in liquid nitrogen and stored at −80°C until mass analysis.

Samples were rapidly thawed and injected onto an ultra-performance liquid chromatography (UPLC) system at 2°C. The protein was run over an immobilized pepsin/anPep column (Affipro) for 2 min at 200 μl/min, and the peptides were collected onto a pre-column trap followed by a 1.5 min desalting wash of the trap column alone (Acquity UPLC® BEH™ C-18 1.7 μm (2.1 x 5 mm)). The trap was eluted in line with a Luna Omega Reverse Phase Column (Polar C18 100 Å 1.6 μm (2.1 mm x 100 mm)) in 0.1% formic acid with a gradient of 10-43% acetonitrile over 16 min at 45 μl/min. Mass spectrometry experiments were performed on an Orbitrap Fusion Lumos (Thermo Scientific) acquiring over a mass range from 300-2,000 m/z, using a standard H-ESI source and the manufacturer’s protocol. Peptides were identified using data-dependent acquisition methods following tandem MS/MS experiments with the top 20 ions selected for fragmentation. MS/MS datasets were analysed using MaxQuant against a database of purified proteins and known contaminants with a false detection rate cut-off of 1%.

Deuterium incorporation of peptides was automatically calculated using HDExaminer Software (Sierra Analytics). Peptides were manually inspected for correct parameters (charge state, retention time, isotopic distribution etc). Results are presented as relative levels of deuterium incorporation with the only control for back exchange as the deuterium levels in the buffer. Changes in any peptide at any time point greater than specified cut-offs (> 5% deuterium incorporation and 0.4 Da between *apo* and different conditions with an unpaired student t-test of p < 0.01) were considered significant.

## Acknowledgements

We gratefully acknowledge funding from the Deutsche Forschungsgemeinschaft DFG (FOR 2848, project TP02 (M.v.d.L.), FOR 2848, project TP06 (O.D.), SFB/TRR 186, Project A23 (O.D.)) and the MDC graduate school (E.N.). The authors gratefully acknowledge Dr. Y. Roske and the entire MX group at BESSY II for support during crystallographic data collection at beamline MX14.1. We are grateful to Dr. A. Schütz (Protein Production and Characterization Platform, MDC) for support with the mass spectrometry validation.

## Contributions

E.N. planned the experiments, designed constructs, solved the crystal structure, performed the biochemical and electron microscopy experiments and assembled the Mic60-Mic19 activation mechanism models. M.R. and A.L. performed the cross-linking experiments and analysed them along with E.N. E.N. and G.D. designed and interpreted the HDX-MS measurements, which were performed by G.D. D.L. performed liposome co-sedimentation and electron microscopy experiments. W.Y.M. generated mutant *S. cerevisiae* strains and performed affinity purification experiments, supervised by K.v.d.M. C.S. and S.K. prepared samples for electron microscopy. A.T. and T.B.B. contributed to the BN-PAGE and SEC-RALS experiments. V.H., M.v.d.L., F.L. and O.D. provided equipment and supervised the research. E.N. and O.D. wrote the manuscript, with inputs from all authors.

## Author information

The atomic coordinates of the extended mitofilin CHCH complex have been deposited in the Protein Data Bank with accession number 28XR. The cross-linking data has been uploaded to the PRIDE server under Project accession number PXD082266 and awaits release. The HDX data raw files can be made available upon request.

